# Epidrug Screening Identifies Type I PRMT Inhibitors as Modulators of Lysosomal Exocytosis and Drug Sensitivity in Cancers

**DOI:** 10.1101/2024.08.26.609671

**Authors:** Baris Sergi, Neslihan Yuksel-Catal, Selahattin Can Ozcan, Hamzah Syed, Umamaheswar Duvvuri, Kirill Kiselyov, Ceyda Acilan

## Abstract

Epigenetic changes drive differential gene expression, contributing to oncogenic transformation and drug resistance. Lysosomes are crucial in cell signaling and the sequestration of toxins and chemotherapeutic agents. This sequestration followed by expulsion through lysosomal exocytosis is a factor in drug resistance. The epigenetic regulation of lysosomal exocytosis remains poorly understood. Our research focuses on this regulation, hypothesizing that epigenetic modifier drugs (epidrugs) capable of inhibiting lysosomal exocytosis and could serve as potential therapeutics. Additionally, we investigate their potential synergy with drugs known to be sequestered in lysosomes.

To examine this concept, we screened approximately 150 epigenetic drugs targeting various reader, writer, or eraser proteins. These drugs were assessed for their combined cytotoxic effects with cisplatin, their impact on lysosomal exocytosis, and on lysosomal biogenesis. Our findings reveal that among the epidrugs showing synergy with cisplatin and further reducing cell viability in combination, two type I PRMT inhibitors, MS023 and GSK3368715, inhibited lysosomal exocytosis. Notably, neither of these drugs altered the expression of the CLEAR lysosomal biogenesis network of genes, suggesting the involvement of novel regulators in lysosomal functions. To explore the specific components of the trafficking machinery affected by PRMT inhibitors, we conducted an RNA-seq analysis, uncovering several differentially expressed genes (DEGs). In addition to previously described functions such as methylation activity, or DNA repair; these DEGs included those involved in vesicular trafficking, lysosomal enzyme activity and lysosome dynamics, offering potential insights into the mechanism of reduced exocytosis and identifying a novel mode for its regulation. Additionally, both inhibitors exhibited synergy with other drugs known to be sequestered in lysosomes, such as carboplatin, oxaliplatin, sunitinib, and doxorubicin, indicating that inhibition of lysosomal exocytosis may be a common phenomenon for such drugs. These findings underscore the potential of Type I PRMT inhibitors as therapeutic agents in cancer treatment. Consistently, analysis on the publicly available patient data revealed that lower levels of type I PRMTs (PRMT1 and 6) were associated with better patient response to these drugs, further suggesting their potential as drug candidates for combination therapy to enhance chemotherapy efficacy and improve cancer patient survival rates.

## Introduction

Tumor resistance to chemotherapy is a growing concern, reducing the effectiveness of cytotoxic drugs across various cancer types and treatments. This resistance, whether innate or acquired, presents significant challenges in drug development and cancer therapy, highlighting the need for innovative strategies [1, 2]. Epigenetic regulation drives gene expression changes that promote oncogenic transformation or drug resistance [3, 4]. Small molecule inhibitors of these epigenetic factors (epidrugs) can induce growth arrest, cell differentiation, or apoptosis [5]. To this end, epidrugs offer a promising solution by altering the epigenetic profile of cancer cells, making them more responsive to conventional therapies [1, 2, 6]. Several epigenetic inhibitors have been FDA-approved for treating various cancers, supported by evidence of their efficacy in adult cardiovascular, neurodegenerative, inflammatory, and specific pediatric lysosomal storage diseases [7–13].

One resistance mechanism in cancer cells is the sequestration of chemotherapeutic drugs within lysosomes [14, 15]. Lysosomes, with their high acidity, degrade and absorb nutrients and damaged cellular components and sequester toxins, along with toxic metals [16–19]. This lysosomal sequestration process is documented for many cytotoxic drugs, including platinum compounds like cisplatin, carboplatin, and oxaliplatin [20, 21]. Especially for organic weak base drugs since the acidity of the lysosomal lumen attracts these drugs naturally [22, 23]. Following sequestration, lysosomal exocytosis expels drugs, where lysosomal membranes fuse with the plasma membrane to release contents into the extracellular space, reducing intracellular chemotherapeutic agent concentrations and diminishing their cytotoxic effects [24, 25]. This process is regulated by molecular motors and cytoskeletal elements, involving complex molecular machinery such as SNAREs, trafficking proteins, cytoskeleton, molecular motors, adaptors, and calcium channels, which remove chemotherapeutic agents and modulate the tumor microenvironment, promoting tumor growth and progression [26, 27]. Lysosomal enzyme release during exocytosis degrades extracellular matrix components, promoting cancer cell invasion and metastasis [28–30].

Previous research revealed that blocking lysosomal exocytosis increases the toxicity and intracellular accumulation of transition metals and platinum compounds [31–34]. Thus, manipulating lysosomal flux has the potential to enhance cancer treatments [26, 32, 35, 36]. Understanding the epigenetic regulators of lysosomal exocytosis could develop interventions that potentiate existing chemotherapeutic agents. Research on epigenetic regulation of lysosomal exocytosis is limited. An epigenetic rheostat, including HDAC2 and TFEB, regulates lysosomal and autophagic pathways [37]. HDAC10 depletion sensitizes cells to doxorubicin by inhibiting drug efflux via lysosomal exocytosis [36]. Reduced expression of Sirtuin 1 (SIRT1) inhibits lysosomal function, promoting breast cancer aggressiveness and survival [38]. These studies highlight the relationship between epigenetic modulation and lysosomal dynamics, suggesting targeted epigenetic interventions could disrupt lysosome-mediated drug resistance mechanisms. Our hypothesis focuses on reversing drug resistance by enhancing the cytotoxic activity of chemotherapeutic agents through inhibiting lysosomal exocytosis with epigenetic regulators.

This study reports a novel mechanism of epigenetic regulation impacting lysosomal exocytosis and its implications for cancer therapeutics. We focused on epidrugs due to the increasingly recognized role of epigenetics in cancer biology and the emerging understanding of lysosomal exocytosis in cancer progression and therapy resistance. Using an epigenetic modifier drug library, we screened for drugs that significantly inhibit lysosomal exocytosis, revealing a poorly understood intersection of these two fundamental biological processes.

## Materials and Methods

### Cell Culture

Du145 (ATCC, HTB-81) cells were maintained in RPMI-1640 (Gibco,11875093), supplemented with 10% FBS (Biowest, S1810) and 1% Pen/Strep (Biowest, L0022) at 37°C and 5% CO2 humidified incubator. Panc1 (ATCC, CRL-1469) cells were maintained in DMEM (Gibco, 11965118), supplemented with 10% FBS and 1% Pen/Strep at 37°C and 5% CO2 humidified incubator. The rest of the cells were maintained at the ATCC’s recommended medium and culturing conditions. Cells were harvested using 0.05% Trypsin-EDTA (Gibco, 25300-054), passaged, or medium refreshed every 3-4 days, depending on the confluency.

### Epigenetic Modifier Drug (Epidrug) Library Screening

The epidrug library containing 148 epigenetic modifier inhibitors was applied (Cayman, 11076) on Du145 cells. For epidrug and cisplatin combination cell viability screening, 1x10^3^ cells were seeded on 96 well plates, and the next day, cells were pre-treated (primed) with epidrugs at a final concentration of 5uM for 72 h. Following priming, cells were subjected to combinatorial treatment with epidrugs and cisplatin at an indicated concentration for a further 72 h (combination). After treatment incubations were done, screenings proceeded to Sulforhodamine B (SRB) cell viability assay, as indicated in the materials and methods section (Figure 2A).

For epidrug library screening with β-Hex lysosomal exocytosis or Lysotracker Green (LTG) staining assays, Du145 cells were seeded on 12 well plates (2x10^5^ cell/well) and 96 well plates (4x10^3^ cell/well), respectively. The next day, cells were treated with epidrugs (at a constant final concentration of 5 μM) for 72 h. Following epidrug treatment, β-Hex lysosomal exocytosis or Lysotracker Green (LTG) staining assays were performed as indicated in the materials and methods section (Figure 2C).

### β-Hex Lysosomal Exocytosis Assay

β-Hex activity was assayed with regular buffer (10 mM HEPES pH 7.4, 150 mM NaCl, 5 mM KCl, 1 mM CaCl2, 1 mM MgCl2, 2 g/L glucose). 2x10^5^ cells were seeded on 12 well plates. The next day, cells were washed once with regular buffer, and cells were incubated with a regular buffer with (stimulated) or without (basal) CuCl2 (Sigma, 222011) for 1 h. Buffer on top of cells was collected and further incubated with 3 mM 4-nitrophenyl-N-acetyl-β-D-glucosaminide substrate (Sigma, N9376) for 1 h at 37 °C. The addition of borate buffer stopped reactions, and the absorbance was measured at 405nm. To determine the total cellular content of β-Hex, cells were lysed with 1% Triton X-100 in PBS, and the cell extract was subjected to the same enzyme activity reaction. Enzyme activity was determined via colorimetric reading as the amount of 4-nitrophenol produced at the end of the reaction, and the activity directly gave the lysosomal exocytosis rate of cells. Absorbance was calibrated with standards made up of 4-nitrophenol (Sigma, N7660) in 0.1M citrate buffer. To determine the total % of β-hex secreted, the following formula was used: β-Hex secreted, % total=[(Abs405 of regular buffer sample)/(Abs405 of regular buffer sample + Abs405 of lysate sample)].

### LysoTracker Green (LTG) Staining Assay

Cellular lysosomal content change along with epidrug treatment was measured by staining with Lysotracker Green dye (Thermo, L7526). The day after seeding Du145 cells at a density of 4x10^3^ cells/well to 96 well plates, drug treatments were performed. At the termination time point, cells were washed with PBS (Biowest, L0615), and medium containing 100 nM LysoTracker Green was added. After incubation for 1 h at 37°C, cells were washed at least 5 times with PBS to get rid of excess dye. Fluorescent measurements were taken using a microplate reader (Synergy H1 Hybrid reader, BioTek). Then, cells were lysed with 25 μl of 0.2% SDS, total protein amount was quantified by BCA assay (Thermo, 23225). Fluorescence values were blank subtracted, and RFU values were normalized to the total protein amount (μg) to get the RFU/μg protein value.

### Sulforhodamine B (SRB) Cell Viability Assay

Prior to drug exposure, cells were seeded in 96 well plates at an indicated density. After treatment incubations, the cells were fixed with a final concentration of 10% (w/v) TCA (Sigma, T6399) for 1 hour at 4°C, washed with deionized water, and allowed to air dry at room temperature. 50 μl of 0.4% (w/v) SRB (Santa Cruz, sc-253615) dye was used to stain the cells for 30 minutes, followed by washing with 1% acetic acid. The SRB dye was dissolved using 150 μl of a 10 mM Tris base solution (Sigma, T1503), and absorbances were measured at 564 nm using a microplate reader (Synergy H1 Hybrid reader, BioTek). Colorimetric measurements were read at 564nm wavelength. Viabilities were calculated as follows; % Viability = [100 x (Sample Abs-blank) / (Non-treated control Abs-blank)].

### Crystal Violet Colony Formation Assay

0.5-2x10^3^ cells were seeded on 12-well plates overnight, and then cells were treated, with the medium being refreshed every 72 hours after treatments were done. Colony formation was expected to occur within 1-3 weeks. After this period, the medium was aspirated, and the wells were washed with PBS (Biowest, L0616). The cells were then fixed with cold methanol for 15- 20 minutes at -20 °C and then washed twice with PBS. Subsequently, the cells were stained with 0.05% (w/v) crystal violet for 30-60 minutes, washed with dH2O, and dried, and the stained colonies were visualized. The percentage of formed colonies was analyzed using ImageJ software.

### Calculation of Combination Index (CI)

The “ComBenefit” software program was used to calculate the combination index (CI) values for two-drug combinations. SRB cell viability assay was used to obtain the dose-response curves for each drug treatment alone and in combination, and the ComBenefit program calculated the synergistic interactions and CI values by the BLISS scoring model. Effect is calculated by [Effect (a+b) = E (a) + E (b) – E (a) E (b)]. A CI value less than 1 indicates synergism, equal to 1 indicates additivity, and greater than 1 indicates antagonism. The results were interpreted according to the program guidelines.

### Western Blotting

Cells were lysed in RIPA buffer (EcoTech, RIPA-100) containing PMSF (Merck, 10837091001) and phosSTOP (Merck, 4906845001) according to the manufacturer’s instructions. Protein concentrations were measured using the BCA Protein Assay Kit (Thermo Fisher, 2322). Proteins were denatured at 95°C for 10 minutes in Laemmli buffer (Bio-Rad, 1610747) with DTT (Sigma, D0632) and separated on SDS-PAGE using Mini-PROTEAN TGX Stain-Free Gels (Bio-Rad, 4568086). Following transfer to PVDF (Bio-Rad, 1620177), membranes were blocked in 5% BSA (Sigma, A3733) and incubated overnight at 4°C with primary antibodies: anti-aDMA (1:5000) (CST, 13522), anti-sDMA (1:2500) (CST, 132228), anti-PRMT1 (1:1000) (Abclonal, A1055), anti-PRMT6 (1:1000) (Abclonal, A5085), and anti-GAPDH (1:5000) (Abcam, AB9485). Membranes were washed with TBS-T, followed by incubation with secondary antibody (Abcam, ab205718) for 1.5 hours at room temperature. Proteins were detected using Immobilon Forte Western HRP substrate (EMD Millipore, WBLUF0500) and visualized on the Licor Odyssey FC Imaging System.

### Real-Time Quantitative PCR (RT-qPCR)

Total RNA was isolated using the NucleoSpin RNA II kit (Zymo, R1051) following the manufacturer’s instructions. 1 µg of total RNA was reverse transcribed using M-MLV Reverse Transcriptase (Invitrogen, 28025-013) cDNA synthesis kit following the manufacturer’s protocol. RT-qPCR was performed using a commercially available LightCycler 480 SYBR Green I master mix (QIAGEN, 1076717) and the designed primers. The reaction mixture included the master mix, cDNA template, and target gene-specific primers (Table 1) in a final volume of 20 μL. Real-time PCR amplification was performed on LightCycler 480 Instrument II (Roche) using the following cycling conditions: initial denaturation at 95°C for 10 minutes, followed by 40 cycles of denaturation at 95°C for 15 seconds, annealing and extension at 72°C for 1 minute. The target gene expressions were normalized to GAPDH housekeeping gene expression, and fold changes were calculated with the 2^(−ΔΔCt) method. To confirm amplification specificity, the PCR products were subjected to a melting curve analysis and subsequent agarose gel electrophoresis if necessary. Statistical analysis was performed using Student’s t-test to compare the expression levels between different groups or conditions with at least three biological replicates.

**Table 1.**
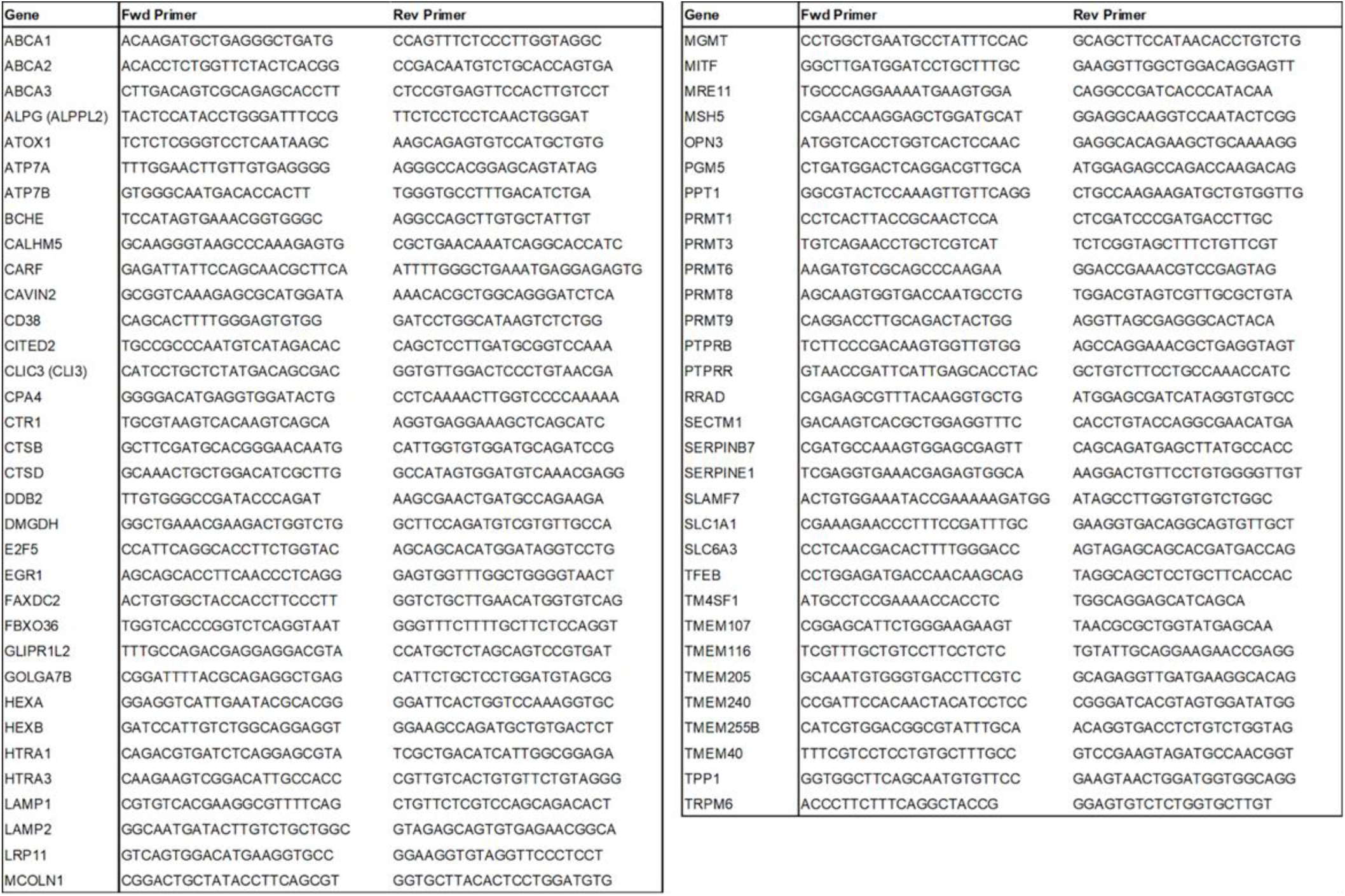
Primers used in the RT-qPCR quantitative gene expression analysis experiments.

### RNA Sequencing and Analysis

Total RNA was isolated with the Qiagen RNeasy Mini Kit following the manufacturer’s instructions. BGI Group conducted poly-A capture, library preparation, and next-generation sequencing as a contract research service. Sequencing was done with BGIseq, which generated 20 million reads/sample and 150bp paired-end reads. The sequencing data was stored securely and analyzed on the Koc University high-performance computing cluster (KUACC). The resulting sequencing data were analyzed with a standard RNA-seq pipeline. First, the quality of the reads was assessed using FastQC software. FastQC provided information on the quality score across reads and the per base sequence content and identified adapter contamination. If contamination was present or there were low-quality regions, then TrimGalore was used for trimming. The sequencing data was then mapped to the most recent human reference genome (NCBI GRCh38) with the software HiSAT2. All mitochondrial, mis-spliced genes and duplicates were identified and removed using RSeQC and Picard Tools. Performing this detailed quality control procedure ensured the success of our study in identifying differentially expressed genes. Once the sequencing reads were aligned, the total read counts for each gene were quantified using featureCounts, part of the Subreads package. The results were then analyzed using the R package DESeq2 to identify differentially expressed genes. The significant results from the gene expression analysis were annotated using the biomart database. The final step was performing Gene Set Enrichment Analysis to understand which pathways or gene networks the differentially expressed genes were implicated in.

## Results

### Cell line panel-based analysis confirms the negative correlation between lysosomal exocytosis and lysosomal volume

Previously published data indicate that chemotherapeutic drugs are sequestered in lysosomes and expelled through lysosomal exocytosis, with the efficiency of this mechanism depending on lysosomal volume and exocytosis rate. Although epigenetic regulation of lysosomal flux is not well understood, the use of epigenetic drug libraries and precise assays presents an opportunity to explore this new translational axis.

To select the optimal cell line for studying lysosomal exocytosis, it was necessary to identify a cell line that yielded satisfactory absorbance values within the linear range for screening changes in exocytosis via β-Hexosaminidase (β-Hex) assay. Additionally, the ideal cell line should exhibit a higher exocytosis rate upon stimulation with copper, which is a common method used to enhance exocytosis by releasing it from lysosomes (**Figure 1A**). After evaluating 19 cell lines, Du145 emerged as the most suitable choice due to its superior performance in the exocytosis assay, alongside Panc1, MIA-PaCa-2, MCF-7, and U373 (**Figure 1B**).

**Figure 1.**
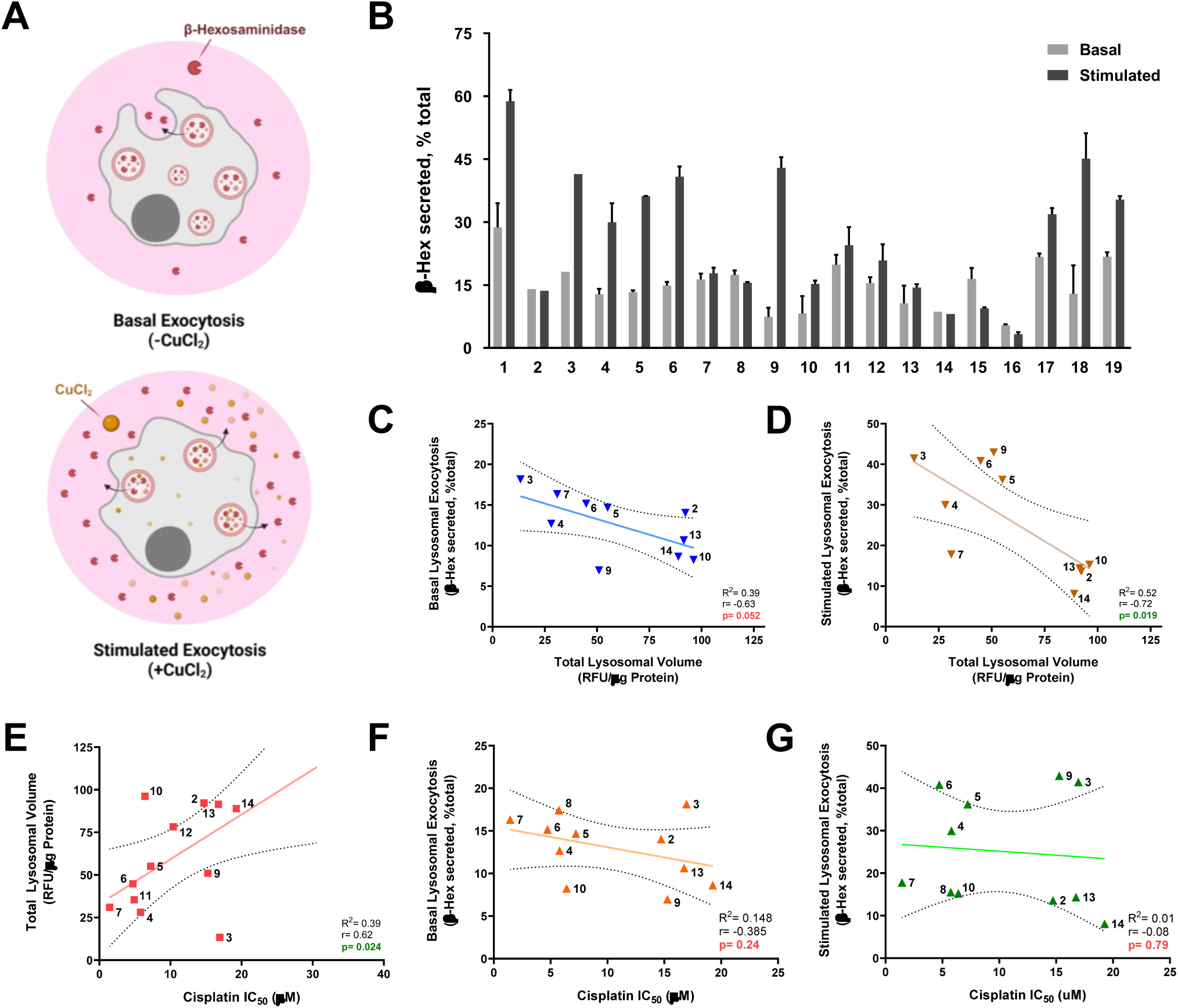
Cell line panel-based studies confirm the correlation between exocytosis and lysosome content. (A) Schematic representation of measuring exocytosis via β-Hex lysosomal exocytosis assay at the basal and CuCl_2_ stimulated conditions (Created with BioRender.com). **(B)** The panel of cell lines was screened for their lysosomal exocytosis rate and induction capacity via CuCl_2_ stimulation. Correlation between total lysosomal volume and lysosomal exocytosis (**(C)** basal, and **(D)** stimulated), **(E)** lysosomal volume and cisplatin IC50, cisplatin IC50 and lysosomal exocytosis (**(F)** basal, and **(G)** stimulated) are plotted. Cell lines were numbered as follows: **1.** MIA-PaCa- 2 (PAAD), **2.** HeLa (CESC), **3.** U2OS (SARC), **4.** U373 (GBM), **5.** PC3 (PRAD), **6.** Du145 (PRAD), **7.** 22Rv1 (PRAD), **8.** LNCaP (PRAD), **9.** H1299 (LUAD), **10.** A549 (LUAD), **11.** HepG2 (LIHC), **12.** Huh7 (LIHC), **13.** MCF-7 (BRCA), **14.** MDA-MB-231 (BRCA), **15.** A2780 (OV), **16.** Kuramochi (HGSOC), **17.** OSCC40 (HNSC), **18.** Panc1 (PAAD), **19.** RPE-1 (Please note that not all 14 cell lines were plotted on specified correlation graphs due to the lack of data for some cell lines. At least 10 cell lines are included for correlation analyses). Error bars represent the SEM of at least three biological replicates (Student’s t-test; *p<0.05, **p<0.01).

In order to gain a deeper understanding of the relationship between lysosomal exocytosis and total lysosomal volume, it is necessary to explore this connection. As inhibiting lysosomal exocytosis leads to a decrease in the release of lysosomal content, we hypothesized that this inhibition would result in an increase in lysosomal volume (size and/or number) within the cell. Our correlation analysis, as shown in **Figure 1C** and **1D**, confirmed the inverse relationship between intracellular acidic vesicle content and exocytosis in both basal and stimulated conditions.

The objective of our study was to investigate whether any of these lysosomal dynamics (exocytosis and total volume) could serve as a predictor of cisplatin sensitivity in cells. We hypothesized that cells with higher exocytosis rates or lysotracker staining would more effectively expel or sequester drugs, leading to reduced cisplatin efficacy and higher IC50 values. Our results demonstrated that while lysotracker staining showed a significant correlation with cisplatin IC50 (**Figure 1E**), no such relationship was observed between exocytosis rates, regardless of whether they were at basal or stimulated levels (**Figure 1F**, **1G**).

### Epigenetic drug library screening on Du145 cells identified epidrugs affecting the lysosomal exocytosis, lysosomal volume, and cell viability in combination with cisplatin

To systematically evaluate epidrugs that enhance the efficacy of cisplatin, with a particular focus on those that may operate through lysosomal sequestration and exocytosis, we conducted three separate screens using a drug library comprising 149 small molecules that target diverse epigenetic ’writer’ (such as Histone Methyl-, Histone Acetyl-, and DNA Methyltransferases), ’eraser’ (including Histone-Deacetylase and -Demethylases), and ’reader’ (containing bromodomains) proteins. It is noteworthy that 33 of these compounds are currently undergoing clinical trials, and 13 have been granted FDA approval. Our approach involved measuring the basal and stimulated exocytosis rates of various cell lines.

The four cell lines with robust activity were evaluated, and the Du145 cells were selected for further screening since their exocytic activity fell within the linear range in both the basal and stimulated states. To assess cytotoxicity in Du145 cells, a mild dose of cisplatin was utilized, ensuring that approximately 70% of the cells remained viable. Additionally, a consistent dose of epigenetic drugs (5 µM) was administered, as these drugs typically lack cytotoxicity and instead prime cells for subsequent therapies by reprogramming cancer cells to a more normal state. Cells were pre-treated with epigenetic drugs for 72 hours and then co-treated with cisplatin for a further 72 hours (**Figure 2A**). **Figure 2B** indicates that the majority of tested epigenetic drugs did not significantly impact cell viability (blue dots, representing each epidrug aligned on the x-axis), suggesting their potential use in sensitizing cancer cells to cisplatin without causing harmful effects on cell viability. A total number of 14 epigenetic drugs that showed decreased cell viability when combined (red dots) were further investigated in cell viability assays with various dose combinations (**Figure 3**).

**Figure 2.**
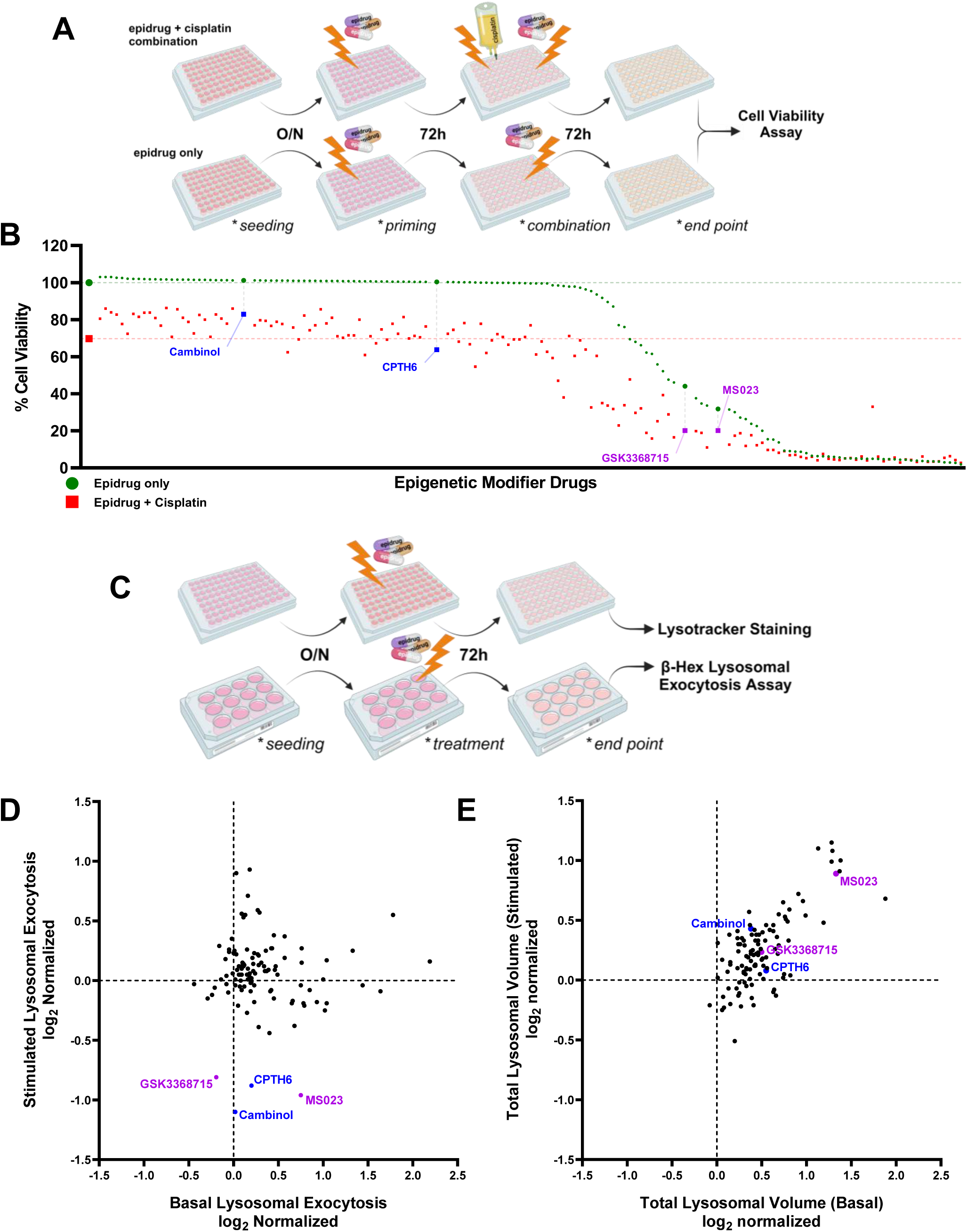
Epidrug library screening identifies epidrugs that synergize with cisplatin and alter lysosomal exocytosis and lysosomal volume in Du145 cells. (A) Schematic representation of epidrug and cisplatin combination cell viability screening (Created with BioRender.com). **(B)** Du145 cells were seeded on 96-well plates and pre-treated (primed) with the epidrugs (5μM) for 72h. Cells were then exposed to 5μM epidrug + 5μM cisplatin combination for another 72h. SRB viability test was conducted at the end of the treatment. Viability was normalized to untreated cells and expressed as % cell viability. Red dashed line corresponds to % cell viability when cisplatin alone is used. **(C)** Schematic representation of lysosomal exocytosis and lysotracker screening experiments (Created with BioRender.com). **(D)** Du145 cells were treated with 148 different epidrugs at 5 μM for 72 h. Exocytosis activity was measured by the extracellular and intracellular activity of β-Hex in basal and stimulated levels. **(E)** Epidrugs were also screened in LysoTracker staining experiments at a constant concentration (5μM) for 72h. The lysotracker signal (RFU/µg protein) in cells in basal and stimulated states are shown after normalization to non-treated control.

**Figure 3.**
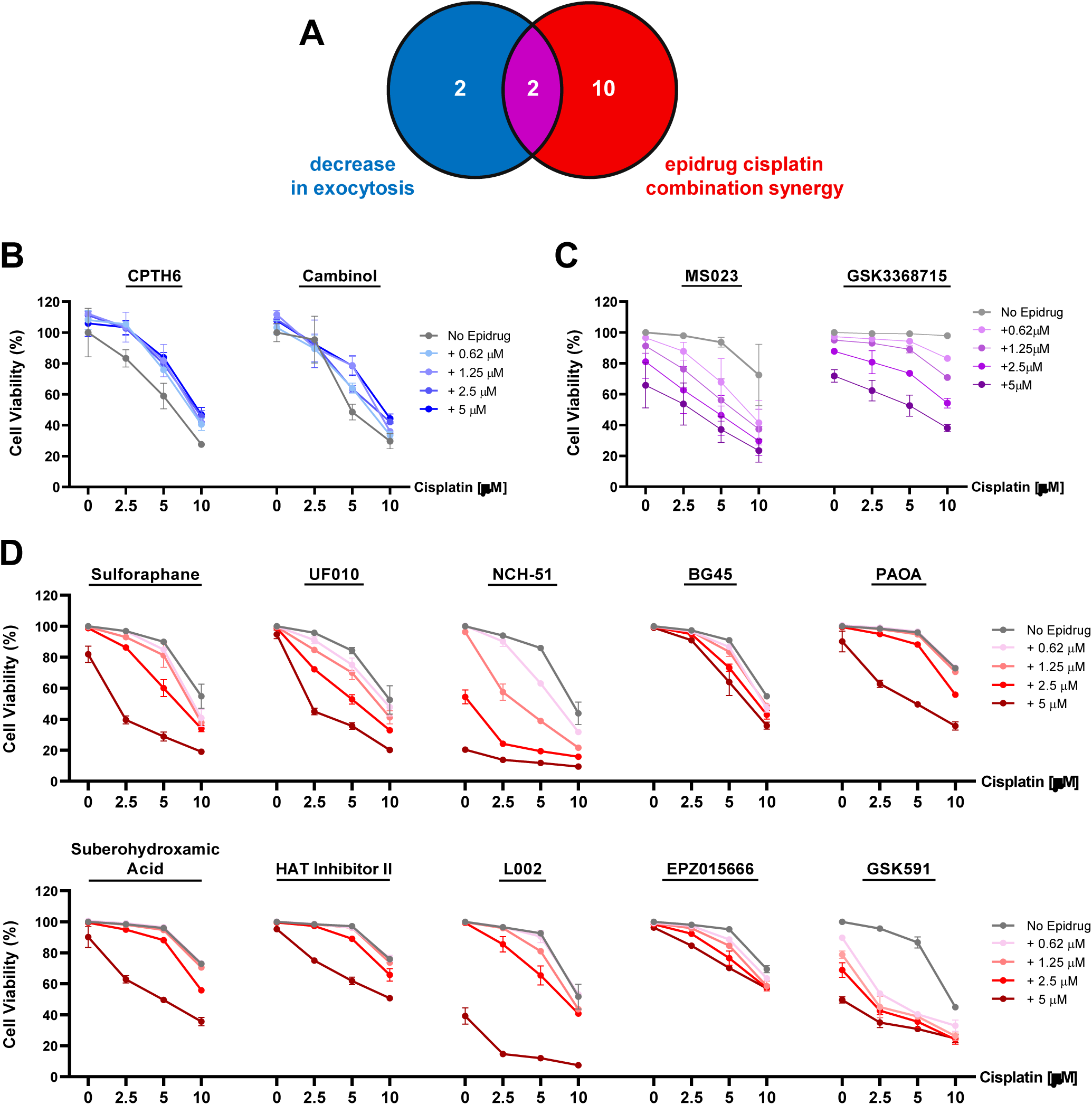
Effect of 14 epidrugs picked from the library on cell viability when combined with cisplatin. (A) Epidrug library screenings were performed on Du145 cells to bring out epidrugs inhibiting lysosomal exocytosis and their potential to synergize with cisplatin in combinatorial treatment regime. In the library, **(B)** 2 epidrugs only decreased lysosomal exocytosis and **(C)** 2 epidrugs both inhibited exocytosis and synergized with cisplatin, **(D)** while 10 epidrugs only synergized with cisplatin.

A comparable methodology was employed for the follow-up screens, in which cells were pre-treated with 5 µM of epidrugs for 72 hours, after which their lysosomal exocytosis and volume were assessed via β-Hex and LysoTracker staining assays, respectively (**Figure 2C**). The motivation for examining lysosomal exocytosis and volume assays in both basal and stimulated conditions was rooted in the hypothesis that we would not diminish the basal level of exocytosis in the cells since the exocytosis activity plays a vital role in fundamental cellular processes. We posited that the inhibition of this lysosomal activity, crucial for numerous cellular processes and whose deficiency would result in lethal consequences, could not be accurately replicated in the experimental setup. As shown in **Figure 2D**, various drugs showed potent and specific effects on lysosomal flux components. Among these, four epidrugs, type I PRMT inhibitors GS3368715 and MS023, a lysine acetyltransferase inhibitor known as CPTH6, and a SIRT1/2 inhibitor called Cambinol, distinguished themselves for their inhibitory effect on lysosomal exocytosis. These four epidrugs also resulted in a considerable accumulation of lysosomal volume (**Figure 2E**), which is consistent with the buildup of lysosomes due to deficiencies in lysosomal exocytosis, as would be expected based on the inverse correlation between exocytosis and lysosome volume (**Figure 1C,D**).

Based on our findings from three separate screens, we have identified 14 potential epidrugs that may influence lysosomal exocytosis, lysosomal volume, and cisplatin efficacy (**Figure 3A**). Through pre-treatment of Du145 cells with these 14 epidrugs in combination with cisplatin at different doses, we discovered that only 2 of the 4 epidrugs that inhibited lysosomal exocytosis also demonstrated synergy with cisplatin (**Figure 3B** and **3C**). The remaining 10 epidrugs, although they did not inhibit exocytosis, did synergize with cisplatin (**Figure 3D**). In detail, 6 of them are HDAC inhibitors, 2 of them are p300 inhibitors, and the remaining 2 are PRMT5 inhibitors. Following these screens, we selected the type I PRMT inhibitors “MS023” and “GSK3368715” for further study as they both inhibit exocytosis and demonstrate synergy with cisplatin.

### Type I PRMT inhibitors MS023 and GSK3368715 alter their target cellular processes on Du145 and Panc1 cells

Protein arginine methyltransferases (PRMTs) are enzymes responsible for the methylation of arginine residues on specific target proteins, a key post-translational modification impacting numerous cellular functions [39]. Type I PRMTs, encompassing PRMT1, -3, -4, -6, and -8, predominantly generate asymmetric dimethylarginine (aDMA) marks on the target protein. This modification plays a significant role in regulating gene expression, signal transduction, RNA processing, and protein-protein interactions [40, 41]. Through the methylation of histones and various nuclear proteins, Type I PRMTs influence chromatin architecture and accessibility, thereby contributing to the epigenetic regulation of gene transcription [39, 42, 43]. Both epidrugs identified as hits in the drug library screens conducted in our study, MS023 and GSK3368715, are defined as drugs that inhibit Type I PRMTs at different inhibition concentrations [44, 45] (**Table 2**). To better understand the properties of these epidrugs, we examined their effects on lysosomal exocytosis, total lysosomal volume, arginine methylation of target proteins, and the expression of lysosomal genes. The epidrugs were tested on two cell lines, Du145 and Panc1, which were among the most suitable cell lines identified in previous experiments (**Figure 1B**). The β-Hex lysosomal exocytosis assay conducted on the Du145 and Panc1 cells showed that the treatment with MS023 and GSK3368715 for 72 hours inhibited lysosomal exocytosis in the stimulated state (**Figure 4A**). Additionally, the ratio of exocytosis at the stimulated and basal conditions (stimulated/basal), which is referred to as induced exocytosis, was reduced post-exposure to epidrugs, indicating that these hit epidrugs alter the way cells release the lysosomal content (**Figure 4B**). Moreover, brief exposure to Type I PRMT inhibitors did not affect lysosomal exocytosis rates, confirming that the reduction in exocytosis required transcriptomic priming of the cells rather than being a direct response to the inhibitor (**Supplementary Figure 1**).

**Figure 4.**
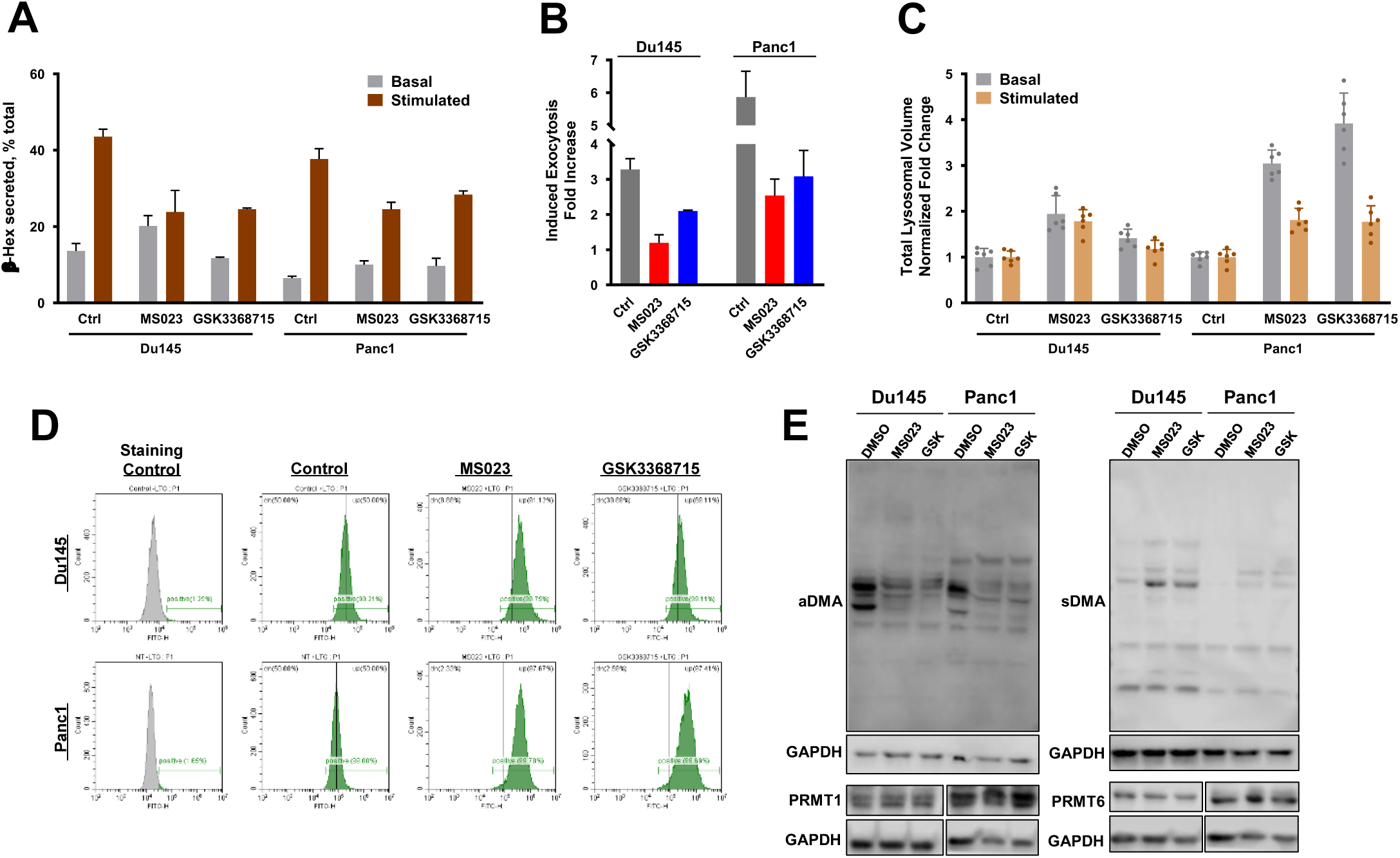
Type I PRMT inhibitors, MS023 and GSK3368715, modulate lysosomal dynamics and change global arginine methylation of target proteins. Du145 and Panc1 cells were treated with MS023 and GSK3368715 (5 uM, 72h), and **(A)** exocytosis was measured at basal and stimulated levels, whereas **(B)** the stimulated exocytosis rates (stimulated/basal) were evaluated as fold increase. Lysotracker staining was performed to determine the total lysosomal volume after epidrug treatment and was shown by **(C)** fluorescence intensity reading and **(D)** flow cytometry. **(E)** Western blot analysis shows the effect of Type I PRMT inhibitors (MS023 and GSK3368715; 5µM, 72 h of treatment) on aDMA and sDMA modification levels, PRMT1, and PRMT6 expressions.

**Table 2.**
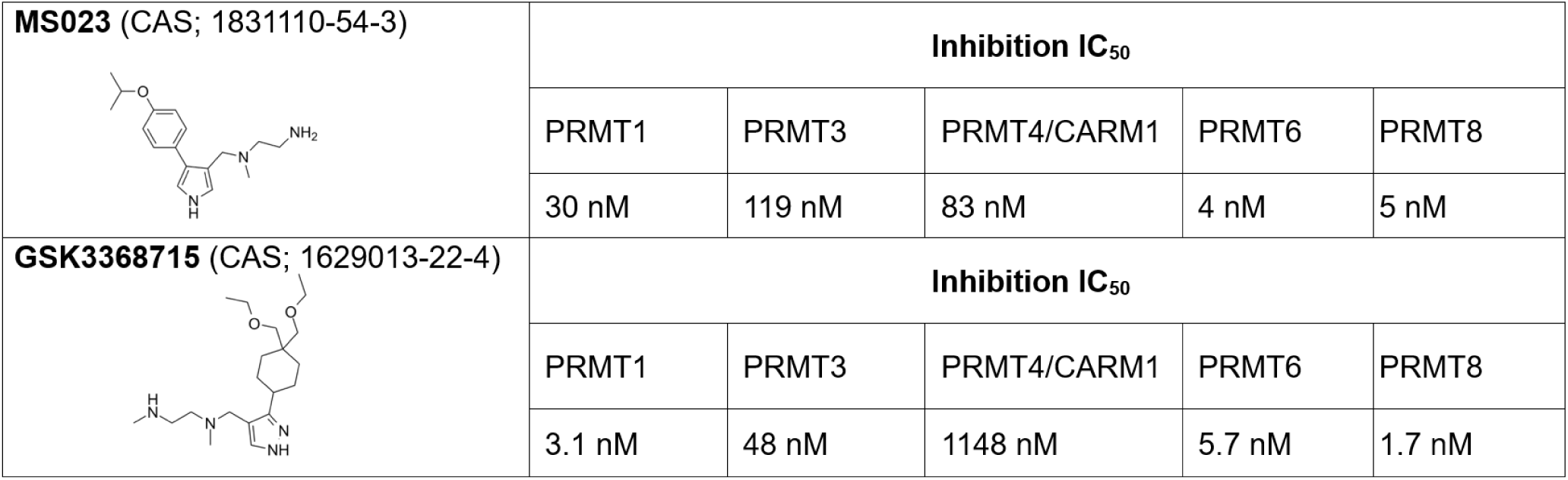
Information about Type I PRMT inhibitors and their inhibition IC50 values on target PRMTs.

| **MS023** (CAS; 1831110-54-3) | Inhibition IC<sub>50</sub> |  |  |  |  |
| --- | --- | --- | --- | --- | --- |
|  | PRMT1 | PRMT3 | PRMT4/CARM1 | PRMT6 | PRMT8 |
|  | 30 nM | 119 nM | 83 nM | 4 nM | 5 nM |
| **GSK3368715** (CAS; 1629013-22-4) | Inhibition IC<sub>50</sub> |  |  |  |  |
|  | PRMT1 | PRMT3 | PRMT4/CARM1 | PRMT6 | PRMT8 |
|  | 3.1 nM | 48 nM | 1148 nM | 5.7 nM | 1.7 nM |

Lysotracker staining assays demonstrated that Type I PRMT inhibitors led to an increase in the total lysosomal volume in Du145 and Panc1 cells (**Figure 4C**). Complementing the fluorescence intensity measurements from Lysotracker staining, flow cytometry experiments showed a rightward shift in the cell population within the treatment groups, further confirming the increase in lysosomal volume (**Figure 4D**).

To assess the impact of the drugs on the stability of type I PRMTs, particularly PRMT1 and PRMT6 -the main targets of MS023 and GSK3368715- we blotted for these proteins. Our findings showed similar levels of protein expression, indicating that the inhibition does not affect protein expression but instead directly influences protein activity (**Figure 4E**). Since type I PRMTs mediate asymmetric dimethylation of arginine residues on target proteins, we also analyzed arginine methylation levels to evaluate target specificity and inhibition efficacy. As expected, treatment with both drugs resulted in decreased levels of asymmetric dimethyl arginine (aDMA) and increased levels of symmetric dimethyl arginine (sDMA, as a compensatory mechanism), confirming effective inhibition of the intended targets (**Figure 4E**).

Lysosomal biogenesis encompasses not only changes in lysosomal volume but also the expression of lysosomal genes. To investigate this, we examined the expression levels of several lysosomal genes, including the master regulators, TFEB and MiTF. Our panel of lysosomal genes included various lysosomal enzymes, transmembrane proteins, ATP pumps, ABC transporters, and proteins involved in lysosomal exocytosis. After treating Du145 and Panc1 cells with MS023 and GSK3368715, we observed that the expression of most lysosomal genes in the panel did not change significantly (**Supplementary Figure 2**). A more detailed analysis revealed that treatment with these inhibitors affected specific genes involved in endolysosomal function, such as TRPML1 (a key regulator of lysosomal exocytosis and biogenesis encoded by MCOLN1 [46]), TMEM205 (a transmembrane protein associated with cisplatin resistance [47, 48]), and ABCA3 (a lipid transporter involved in lysosomal function [49, 50]). Interestingly, while TRPML1 is part of the CLEAR lysosomal biogenesis network, MS023 did not broadly impact other components of this network, including the structural protein LAMP1, the enzyme palmitoyl-protein thioesterase 1 (PPT1), or genes encoding various cathepsin lysosomal proteases (**Supplementary Figure 2A, B**). We conclude that type I PRMT inhibitors selectively affect components of the trafficking machinery related to lysosomal exocytosis without broadly disrupting the entire lysosomal biogenesis network.

We further investigated whether epidrugs exhibit synergistic effects with other chemotherapeutic agents, beyond cisplatin, that are known to undergo lysosomal sequestration and exocytosis, which limits their efficacy. Sunitinib, a receptor tyrosine kinase inhibitor, and Doxorubicin, a topoisomerase II inhibitor, are among the drugs that experience lysosomal sequestration [51–54]. We combined the inhibitors MS023 and GSK3368715 with Sunitinib, Doxorubicin, and platinum-based drugs such as cisplatin, carboplatin, and oxaliplatin on Du145 and Panc1 cell lines. The Combenefit analysis demonstrated that both epidrugs exhibited synergistic effects with these chemotherapeutic agents across various concentrations in both cell lines (**Figure 5**). The colony formation assay further confirmed this synergy by showing a reduction in the cells’ colony-forming capacity after the combination treatment (**Figure 5C**). These results suggest that the inhibition of lysosomal exocytosis by epidrugs enhances the cytotoxicity of the chemotherapeutic agents. Notably, similar to the need for reduced exocytosis rates (**Supplemental Figure 1**), the synergy between epidrugs and agents like cisplatin and sunitinib was only observed when cells were pre-treated with the epidrugs for 72 hours; co-treatment without this priming did not produce any synergistic effects (**Supplementary Figure 3**).

**Figure 5.**
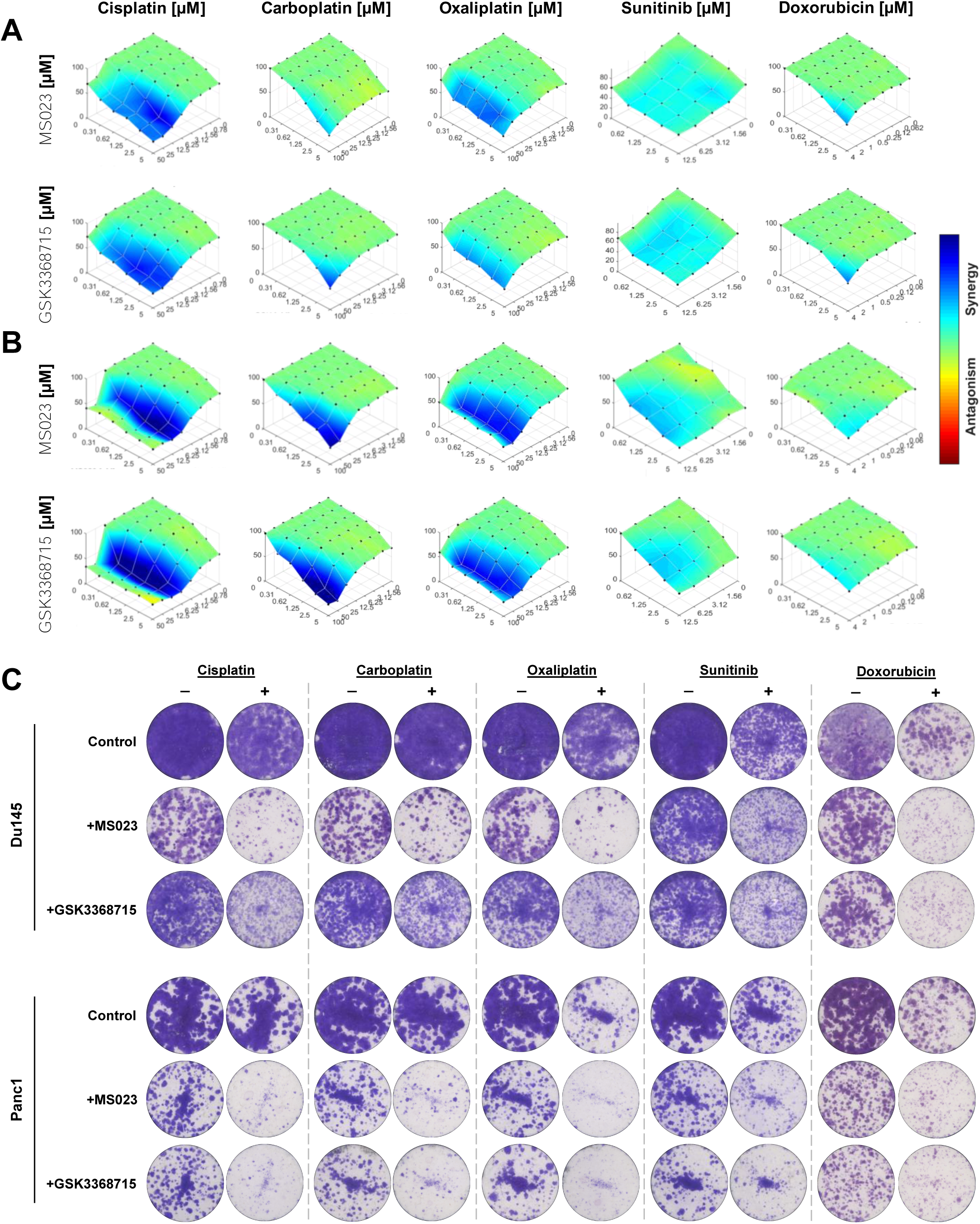
Type I PRMT inhibitors synergize with platinum drugs and chemotherapeutic agents known to undergo lysosomal sequestration. (A) Du145 cells and **(B)** Panc1 cells were treated with MS023 and GSK3368715 in combination with cisplatin, carboplatin, oxaliplatin, sunitinib, and doxorubicin at the indicated doses. Cells were primed with epidrug for 72 h, then combination treatments were carried out with indicated drugs for 72 h. The evaluation of synergistic effects was carried out utilizing Combenefit employing the HSA model, where the deepest shade of blue indicates the highest level of synergy. **(C)** Colony formation assay performed on Du145 and Panc1 cells also confirmed the synergistic effect through inhibition of colony forming capacity of cells after the combination of each epidrug with chemotherapeutic agents.

### RNA Sequencing Analysis on MS023 Treated Cells Reveal Differentially Expressed Genes and Enriched Gene Sets

To explore the synergistic effects of type I PRMT inhibitors with chemotherapy agents and their influence on lysosome volume and exocytosis, we performed RNA-seq analysis on Du145 cells treated with MS023. The results, presented in **Figure 7A** as a volcano plot, show differentially expressed genes (DEGs) that were further examined. Additionally, the expression changes of the top up- and down-regulated genes within the DEGs were validated through qPCR experiments (**Supplementary Figure 4**). Gene ontology analysis confirmed the previously known roles of PRMTs, such as their involvement in DNA repair, methylation, and epigenetic modifications (**Figure 6A**), supporting the robustness of our experimental approach. The analysis also revealed the effects of MS023 treatment on lysosomal flux, showing a negative enrichment in genes associated with lysosomal hydrolases, lipases, peptidases, and catalytic activity (**Figure 6A, B**). Moreover, GO analysis identified significant changes in cellular components, particularly in vesicular transport, coated vesicles, and vesicle membranes (**Figure 6C**).

**Figure 6.**
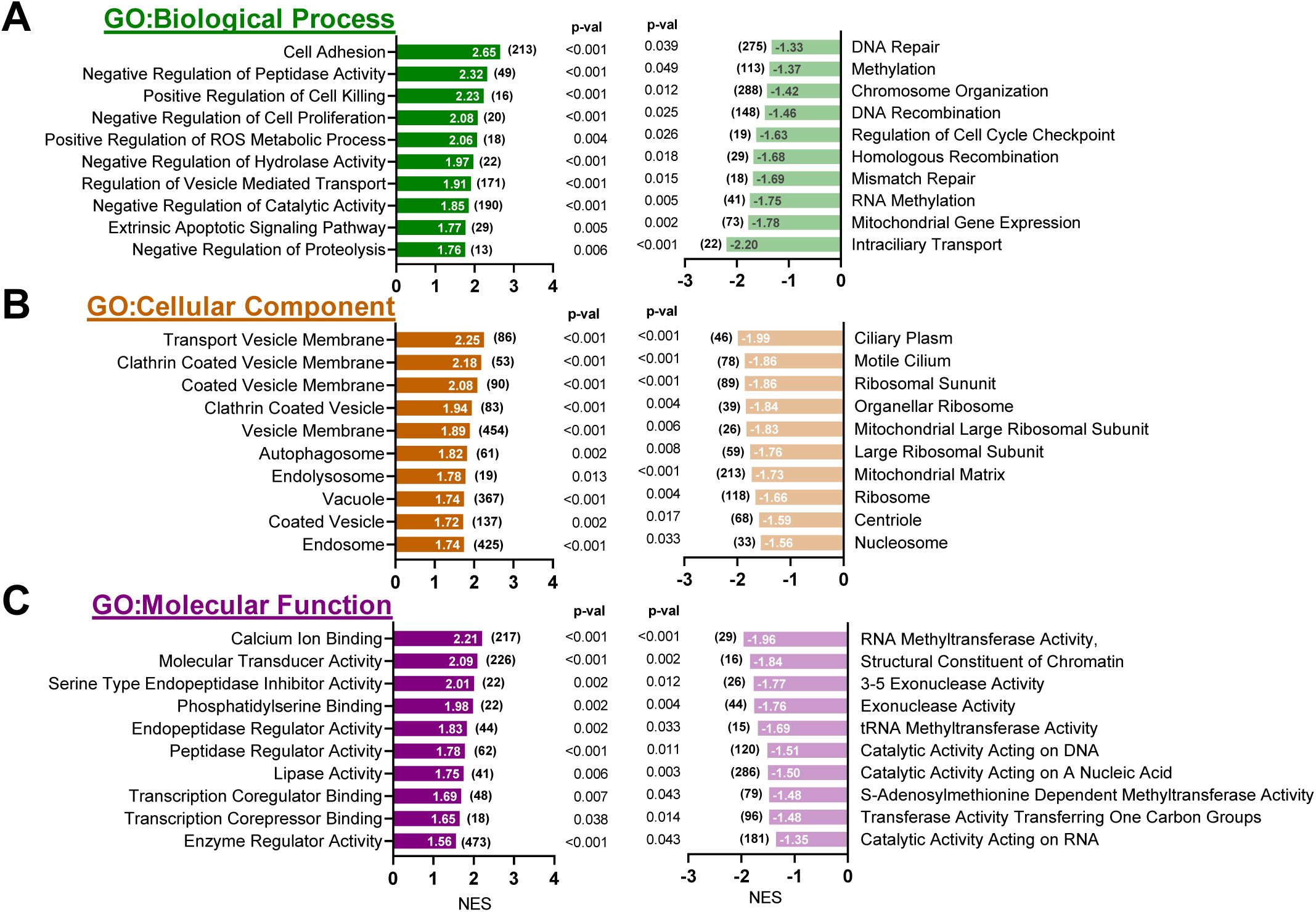
RNA-seq analysis confirms the enriched pathways and gene sets after MS023 treatment in Du145 cells. GSEA analysis was performed using Gene Ontology (GO) gene sets to reveal positively (left panel) and negatively (right panel) enriched gene sets. The numbers in the columns indicate the NES scores, whereas the numbers next to the columns in brackets indicate the size of enriched gene numbers. NOM p-val values are shown next to the columns. GSEA was performed using **(A)** Gene Ontology, Biological Process “c5.go.bp.v2023.1.Hs.symbols.gmt”, **(B**) Gene Ontology, Cellular Components “c5.go.cc.v2023.1.Hs.symbols.gmt”, and **(C)** Gene Ontology, Molecular Function “c5.go.mf.v2023.1.Hs.symbols.gmt” datasets in the MsigDB database.

**Figure 7.**
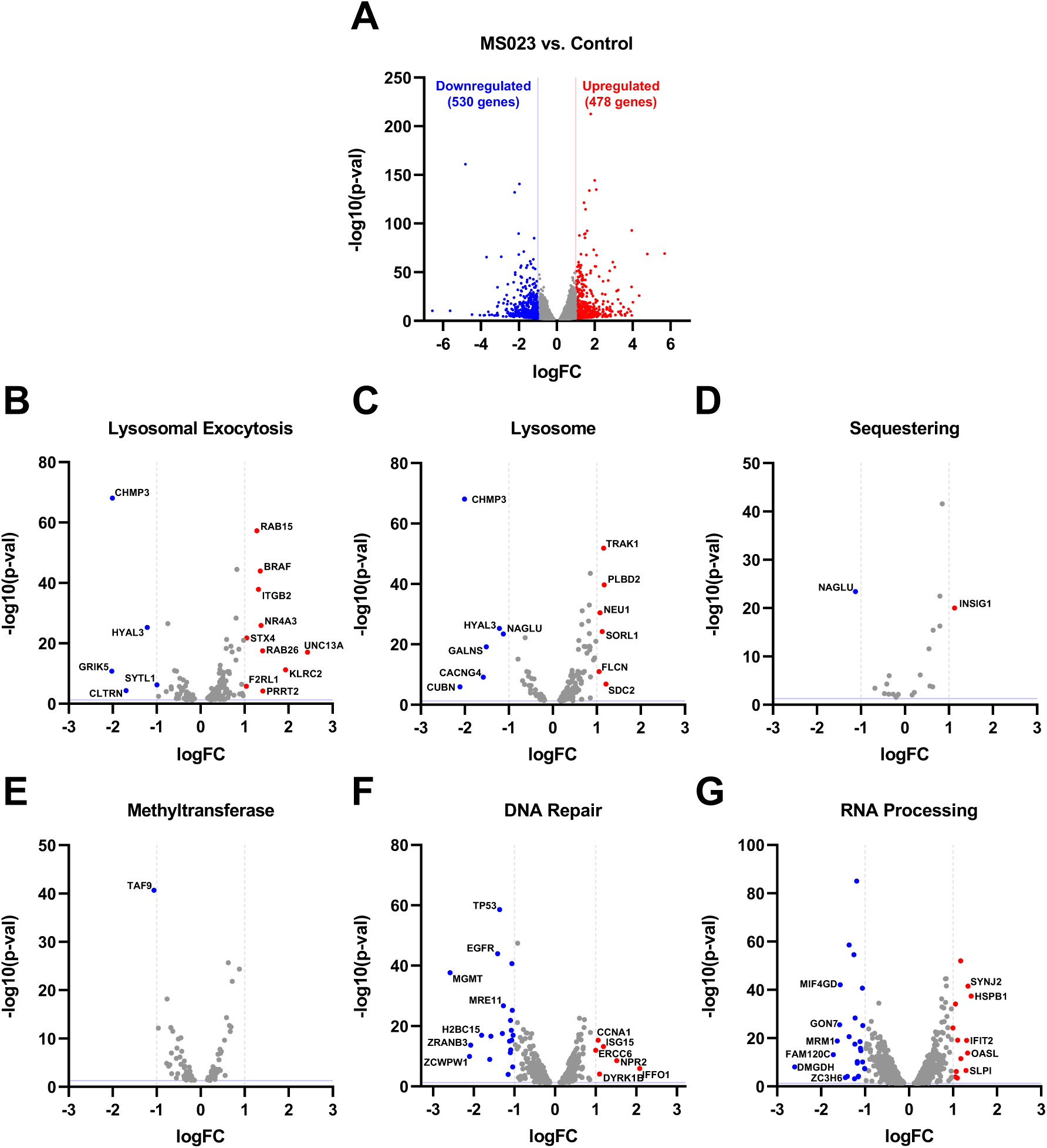
RNA-seq analysis identifies differentially expressed genes within combined gene sets. (**A**) The volcano plot from the RNA-seq analysis shows the differentially expressed genes following MS023 treatment (p-val < 0.05; |log(FC)| > 1). Gene sets from the GSEA Molecular Signature Database were manually selected and categorized into six distinct “combined gene sets” based on their functional relevance (details given in Table 3): (**B**) Lysosomal exocytosis-related gene sets, (**C**) Lysosome-related gene sets, (**D**) Sequestration-related gene sets, (**E**) Methyltransferase-related gene sets, (**F**) DNA repair-related gene sets, and (**G**) RNA processing-related gene sets. The differentially expressed genes within each of these combined gene sets are marked, and volcano plots were generated, with grey vertical lines indicating the |logFC| > 1 cutoff and the blue horizontal line representing p-val < 0.05.

**Table 3.** The list of gene sets being used to curate combined gene sets for RNA-seq analysis.

|  |
| --- |
| **Exocytosis-related Gene Sets**<br>GO:0006887, GO:0045955, GO:2000301, GO:0045956, GO:0045055, GO:0017158, GO:0017157, GO:2000300, GO:0016079 |
| **Lysosome-related Gene Sets**<br>GO:0008333, GO:0032510, GO:0090160, GO:0006622, GO:1905671, GO:0062196, GO:0150031, GO:0004563, GO:0015929, GO:0007042, GO:0097212, GO:1905146, GO:0007041, GO:0035751, GO:1905165, GO:0033299, GO:0098574, GO:0043202, GO:0097401, GO:0007035, HP:0004356 |
| **Sequestering-related Gene Sets**<br>GO:0051238, GO:0140487, GO:0140313, GO:0140311 |
| **Methyltransferase-related Gene Sets**<br>GO:0035242, GO:0008276, GO:0008169, GO:0034708, GO:0035097, GO:0140939 |
| **DNA Repair-related Gene Sets**<br>GO:0006281, GO:0006302, GO:0036297, GO:0006289, GO:1990391, WP4946, WP4753, R-HSA-5693532, R-HSA-73894, R-HSA-5696398, hsa03420, Pubmed 26771021, Pubmed 17891185 |
| **RNA Processing-related Gene Sets**<br>GO:0008380, GO:0001510, GO:0009451, GO:0003723 |

Notably, there was an enrichment of autophagosomes, endolysosomes, and vacuoles following MS023 treatment, aligning with its impact on lysosomal exocytosis and the observed increase in lysosomal content (**Figure 6B**). The enrichment of coated vesicles, particularly in clathrin-coated vesicle formation (NES scores >2), speculatively suggests increased endocytosis and transport of coated particles from the trans-Golgi network to the lysosomes, contributing to the observed increase in lysosomal content (**Figure 2E, 4C**).

In addition to GSEA analyses, we examined existing gene sets in the Molecular Signature Database. Based on the intracellular effects of MS023 reported in the literature and its observed impact on lysosomal functions, we generated six main combined gene sets: Exocytosis-, Lysosome-, Sequestering-, Methyltransferase-, DNA Repair-, and RNA Processing-related combined gene sets. **Table 3** provides a summary of the composition of each combined dataset. We then overlapped the DEGs from our RNA-seq analysis with these combined gene sets to determine which DEGs might be involved in these functions, potentially explaining the reduction in lysosomal exocytosis or the observed drug synergy. Indeed, several genes were identified, and the top up-regulated and down-regulated genes (|logFC| > 1) are highlighted in the volcano plots (**Figure 7**).

We then retrieved ChIP-seq data from public datasets on the ChIP Atlas platform and examined the occupancy of type I PRMTs (namely PRMT 1, 4 and 6 based on available data) at the promoter regions of DEGs identified in the combined gene sets generated in **Figure 7**. This analysis focused on genes with significant upregulation or downregulation in our RNA-seq data (|logFC| > 1) to better understand how PRMT-mediated methyltransferase activity at these promoter regions influences gene expression changes (**Figure 8A**). The presence of type I PRMT binding sites at the promoter regions of target genes, along with the significance of their occupancy, is visualized in a bubble graph, with check marks indicating which gene set the DEG was detected in (**Figure 8B, C**). Consistent with the role of PRMTs in regulating the expression of these genes, we observed type I PRMT occupancy at the promoter regions of several DEGs. These observations strongly support the hypothesis that type I PRMTs directly regulate the expression of specific genes involved in lysosomal functions and related pathways, highlighting the potential of PRMT inhibitors like MS023 to modulate epigenetic marks and alter gene expression, ultimately impacting cellular processes such as lysosomal exocytosis.

**Figure 8.**
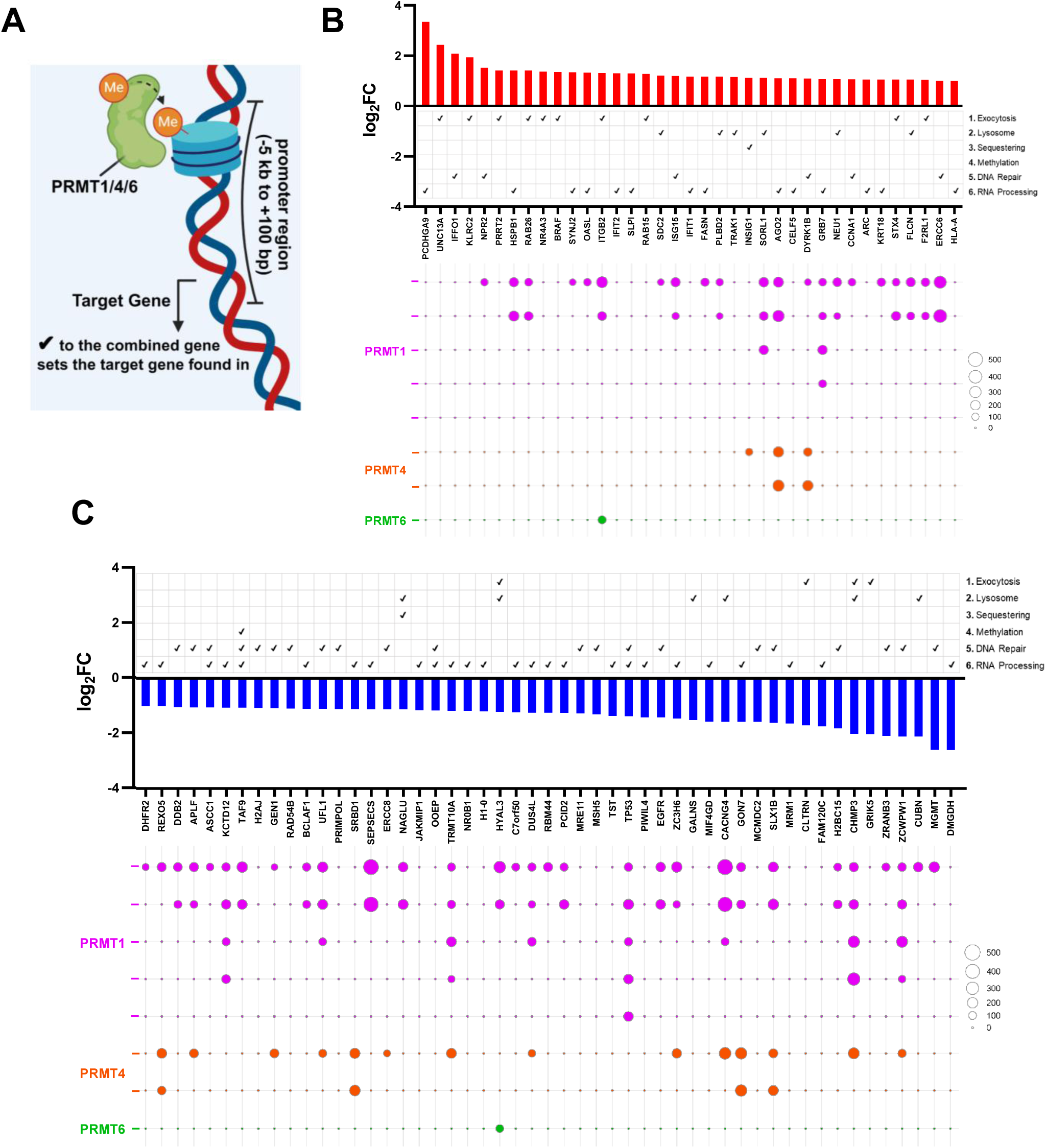
Integrative analysis using RNA-seq data and online available ChIP-seq data reveals the promoter regions of differentially expressed genes found in combined gene sets are occupied by Type I PRMTs. (A) Schematic illustration of the data presented in the figure. Promoter regions of the target genes (-5 kb to +100 bp) are monitored whether PRMT1, -4, or -6 has an enrichment peak in ChIP-seq data retrieved from ChIP Atlas platform (Created with BioRender.com). LogFC values of **(B**) upregulated and **(C)** downregulated genes in combined gene sets (combined gene sets were ticked for a particular target gene) were shown as bar graph, along with the bubble graph indicating enrichment peaks and enrichment scores of PRMTs at the promoter region. In the bubble graph, each line representing PRMT1 (purple), PRMT4 (orange), and PRMT6 (green) corresponds to a distinct enriched peaks on the promoter of target genes, derived either from the same ChIP-seq dataset or from different ones.

To further validate our findings and assess the clinical relevance of PRMT-1, -4, -6, and -8 in chemotherapy resistance, we analyzed patient data from the Cancer Treatment Response Gene Signature Database (CTR-DB), a publicly available resource. This analysis aimed to determine whether the expression levels of type I PRMTs could serve as predictive biomarkers for chemotherapy response, particularly for treatments known to involve lysosomal sequestration. We specifically compared the expression levels of these PRMTs between patients who responded to therapy and those who did not, to establish a connection between our in vitro results and actual patient outcomes.

The patient cohort consisted of individuals diagnosed with a specific type of cancer, divided into two groups based on their response to standard chemotherapy. Responders were defined as patients who exhibited a significant reduction in tumor size or disease stabilization, while non-responders showed no improvement or disease progression despite treatment during treatment [55]. Using this classification, we focused on the patient groups who take specific chemotherapy drugs which attributed to the lysosomal sequestration and exocytosis events, including cisplatin [34, 56], carboplatin [57], Imatinib [58, 59], Sorafenib [60], and Bevacizumab [61]. Our analysis did not find any significant association between PRMT-3, -4, or -8 expression and chemotherapy resistance. However, we observed that PRMT1 and PRMT6 expression levels were significantly higher in the non-responder group compared to the responder group under the specified treatment regimens, consistent with our hypothesis (**Figure 9**). The correlation between low PRMT1 and PRMT6 expression and better therapeutic response reinforces our in vitro findings, supporting the idea that inhibiting type I PRMTs can reduce lysosomal exocytosis-mediated drug tolerance and enhance chemotherapy efficacy.

**Figure 9.**
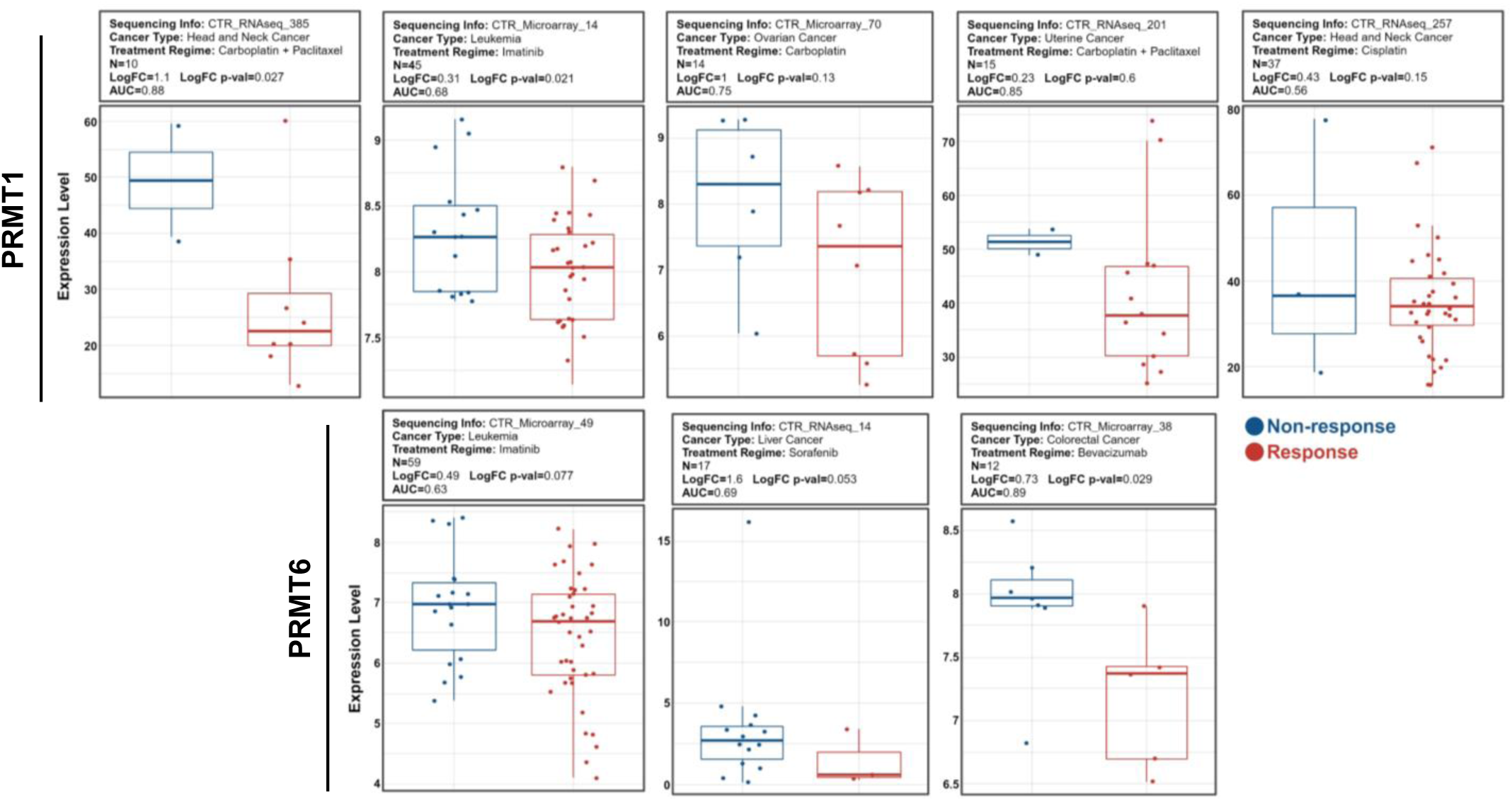
Patient data confirm the contribution of PRMT1 and PRMT6 expressions to the response of patients to chemotherapy treatment regimes. In cohorts of cancer patients exhibiting negative response (non-response) to the indicated treatment regime, relatively higher expression levels of PRMT1 or PRMT6 are observed compared to the responsive patients. Data was obtained from the Cancer Treatment Response Gene Signature Data Base (CTR-DB).

## Discussion

This study aimed to explore the intersection of epigenetic regulation and lysosomal exocytosis in cancer therapy, particularly in overcoming chemotherapeutic resistance. Our findings underscore the significance of lysosomes in sequestering and expelling chemotherapeutic drugs, a process that contributes to drug resistance. By targeting the epigenetic regulation of lysosomal exocytosis, we sought to enhance the therapeutic efficacy of chemotherapy.

We implemented a comprehensive screening process that utilized an epigenetic drug library in order to pinpoint epidrugs that modulate lysosomal exocytosis. The identification of specific epidrugs that inhibit lysosomal exocytosis without broadly affecting lysosomal biogenesis genes underscores the specificity and clinical utility of our findings. We narrowed our analysis to two epidrugs, MS023 and GSK3368715, which target the same epigenetic modifiers, type I PRMTs, emerging as potential molecules that synergize with cisplatin while reducing exocytosis. Notably, the efficacy of the drug combination varied depending on the chemotherapeutic agent used. RNA-seq analysis revealed that these epidrugs affected genes involved in several pathways, including lysosomal function and DNA repair. This suggests that drugs like cisplatin, which cause DNA damage and are sequestered to lysosomes, can synergize through both mechanisms, while Sunitinib, acting through a different pathway, shows less synergy, potentially due to its reliance solely on reduced exocytosis. The observed synergy highlights the potential of using epigenetic drugs to enhance the effectiveness of DNA-damaging agents in cancer therapy by simultaneously targeting multiple pathways, including lysosomal exocytosis.

Type I PRMTs regulate critical cellular processes, including DNA damage response, repair, RNA processing, and splicing [62, 63]. PRMT1 and PRMT6 influence DNA repair by methylating key proteins [64, 65], while PRMT4 modulates RNA splicing under stress [66]. PRMT8, primarily studied in the nervous system, is involved in RNA processing [67]. Targeting these enzymes with inhibitors is a promising therapeutic strategy, particularly in cancer. Inhibition of PRMT1 in triple-negative breast cancer (TNBC) cells triggers an interferon response, leading to cell death [68], and disrupts chromatin recruitment, enhancing cisplatin efficacy [69]. Our results indicate that, in addition to the previously discussed mechanisms, Type I PRMT inhibitors decrease lysosomal exocytosis—a novel finding that, to our knowledge, is being described for the first time. This discovery further enhances the potential of these inhibitors in improving chemotherapy efficacy.

Building on our RNA-seq analysis and the integration of online ChIP-seq data, we propose that MS023 treatment alters histone methylation patterns by inhibiting PRMT activity, thereby affecting the transcriptional regulation of target genes, including lysosomal genes. Our findings demonstrate how type I PRMT inhibitors can modulate gene expression at the transcriptional level, which aligns with existing literature. Moreover, the correlation between low PRMT1 and PRMT6 expression and better therapy response in patient data underscores the relevance of our in vitro results. This supports the idea that type I PRMT inhibitors can mitigate lysosomal exocytosis-mediated drug resistance, potentially enhancing the effectiveness of chemotherapeutic agents, as shown in our cell line studies.

The discovery of MS023 and GSK3368715 as promising candidates for combination therapy, identified through a comprehensive epigenetic drug library, opens new avenues for epigenetic drug research in cancer treatment. However, several key questions remain. The precise mechanisms by which these and other epigenetic drugs influence lysosomal exocytosis and biogenesis need further exploration. Additionally, identifying the specific cancer types that may benefit from this synergistic effect is crucial for advancing these findings into clinical applications. Nevertheless, our results pave the way for innovative therapeutic strategies that hold significant potential to improve cancer treatment outcomes.

## Acknowledgements

The authors gratefully acknowledge the use of the services and facilities of the Koc University Research Center for Translational Medicine (KUTTAM), funded by the Presidency of Turkey, Head of Strategy and Budget. The following figures were created using BioRender.com and licensed for publication (**Figure 1**: OP278AR62J, **Figure 2**: DK278ARE9Z, **Figure 8**: YW278ARJPF)

---

**Supplementary Figure 1.**
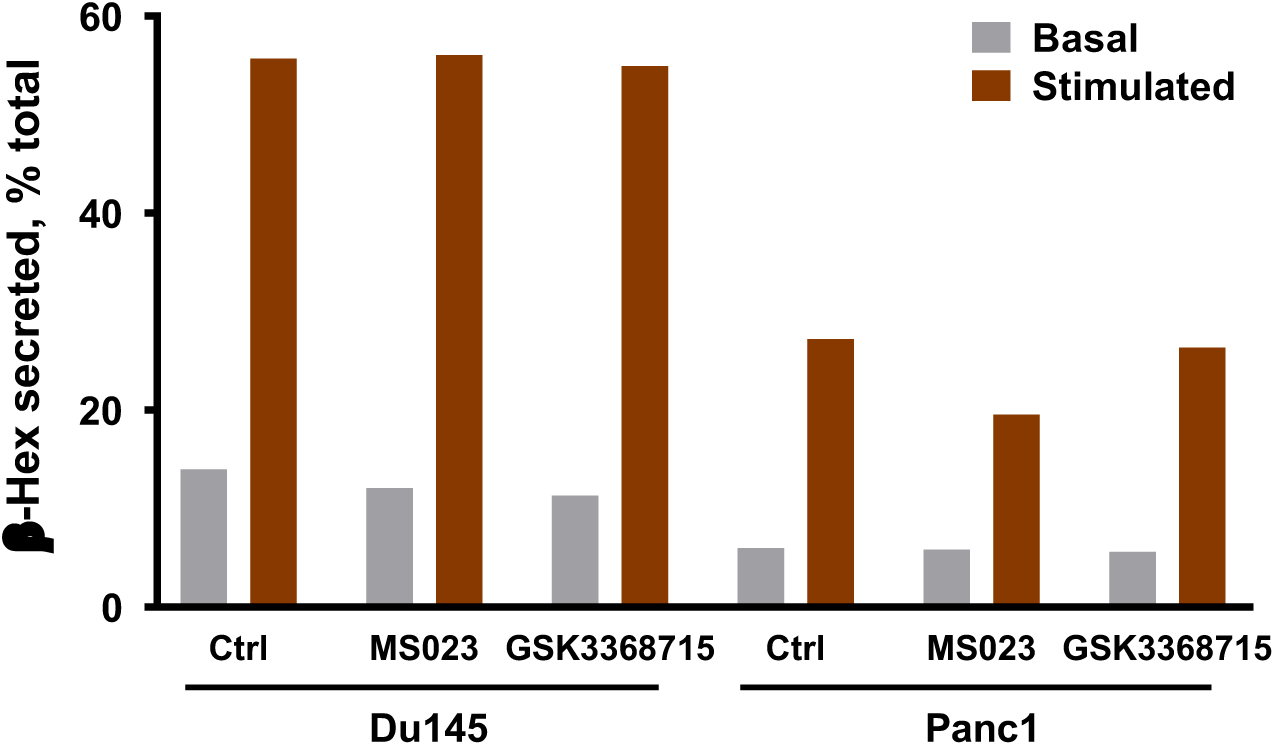
Type I PRMT inhibitors MS023 and GSK3368715 do not inhibit the lysosomal exocytosis upon brief exposure. Du145 and Panc1 cells were briefly exposed (1 h) to MS023 and GSK3368715 epidrugs, then subjected to β-Hex lysosomal exocytosis assay with and without stimulation with CuCl_2_.

**Supplementary Figure 2.**
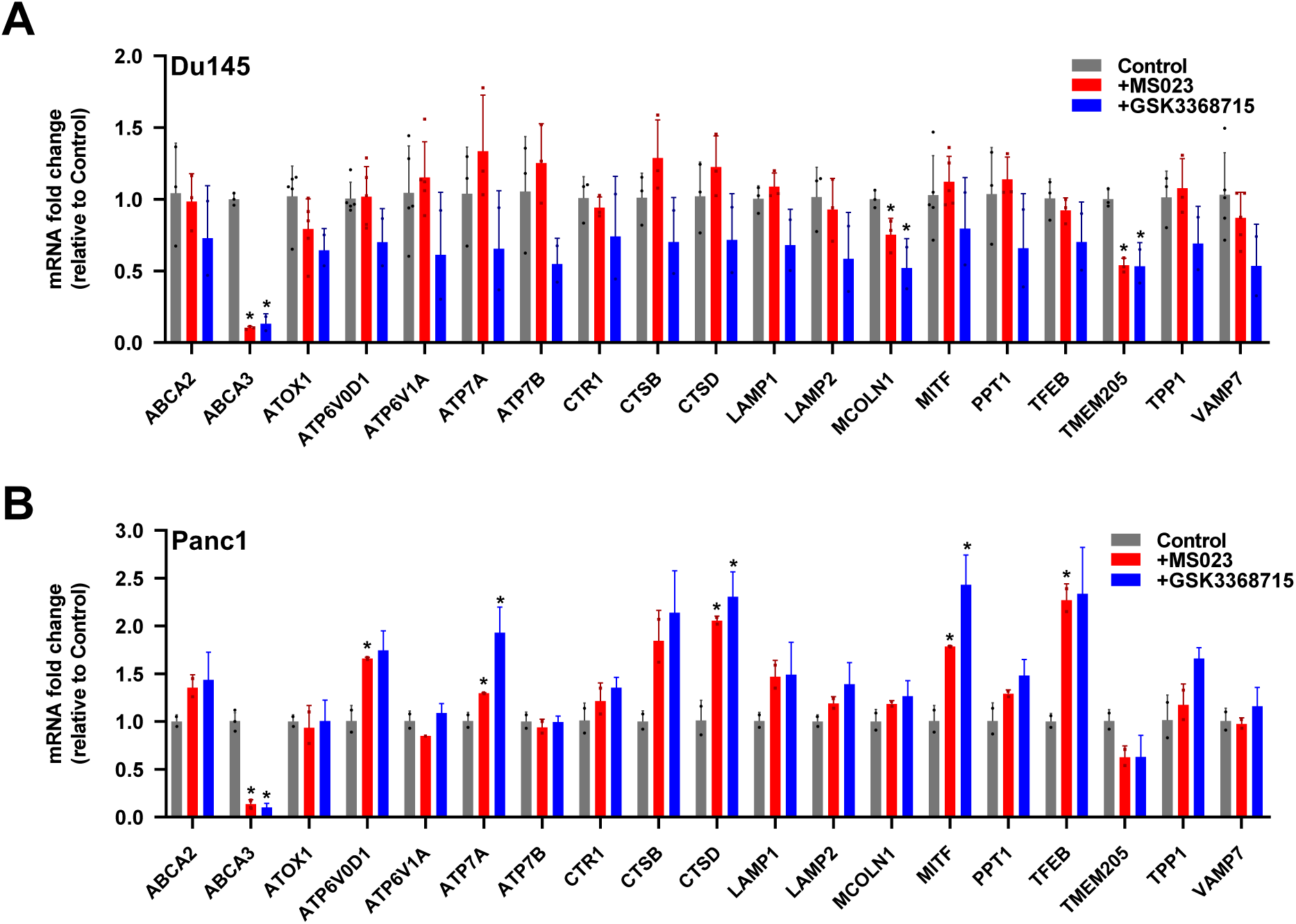
Type I PRMT inhibitors, MS023 and GSK3368715, do not cause an overall expression change in lysosomal gene network. Expressions of genes in the lysosomal gene panel were quantified after MS023 and GSK3368715 treatments (5 uM, 72 h) in **(A)** Du145 and **(B)** Panc1 cells.

**Supplementary Figure 3.**
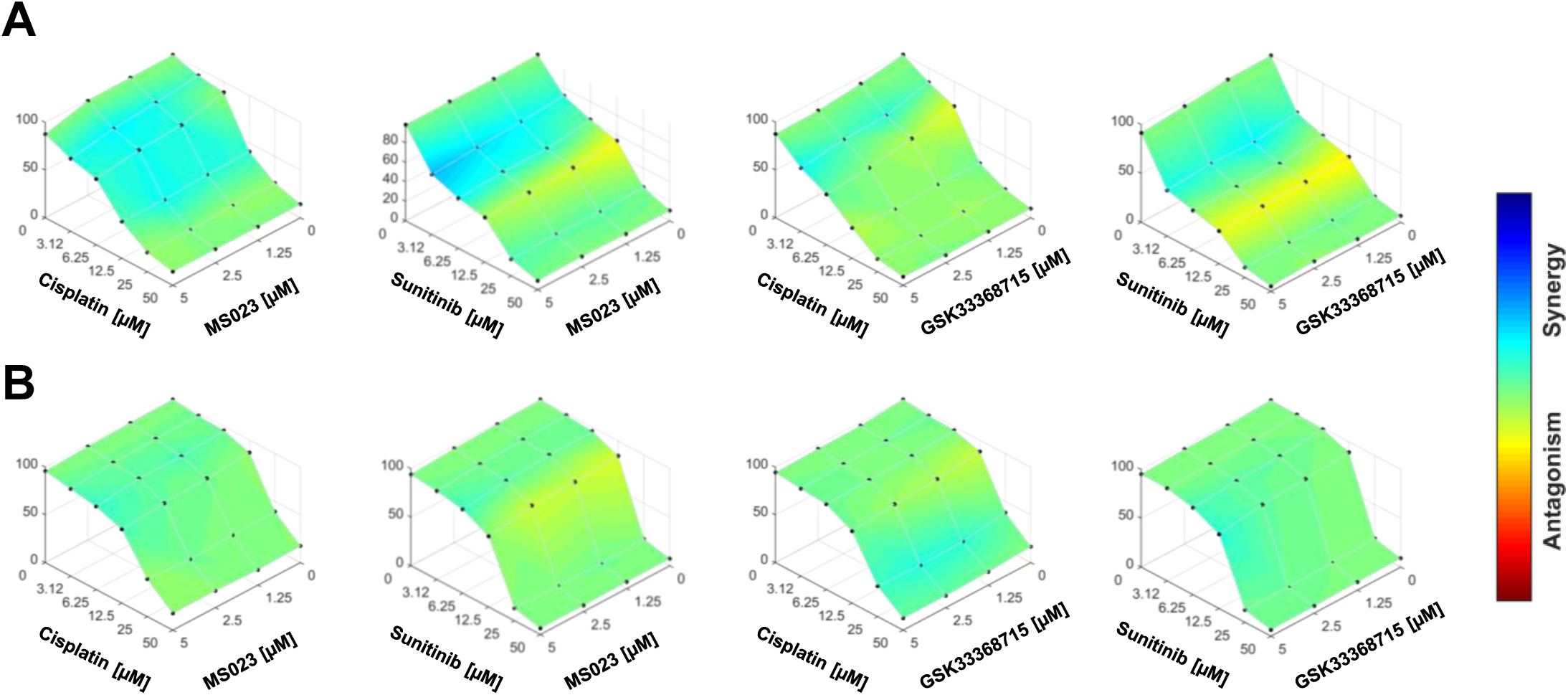
Type I PRMT inhibitors MS023 and GSK3368715 do not synergize with chemotherapeutic agents without priming. Epidrugs were only co-treated with cisplatin or sunitinib for 72 h without priming on **(A)** Du145 and **(B)** Panc1 cells.

**Supplementary Figure 4.**
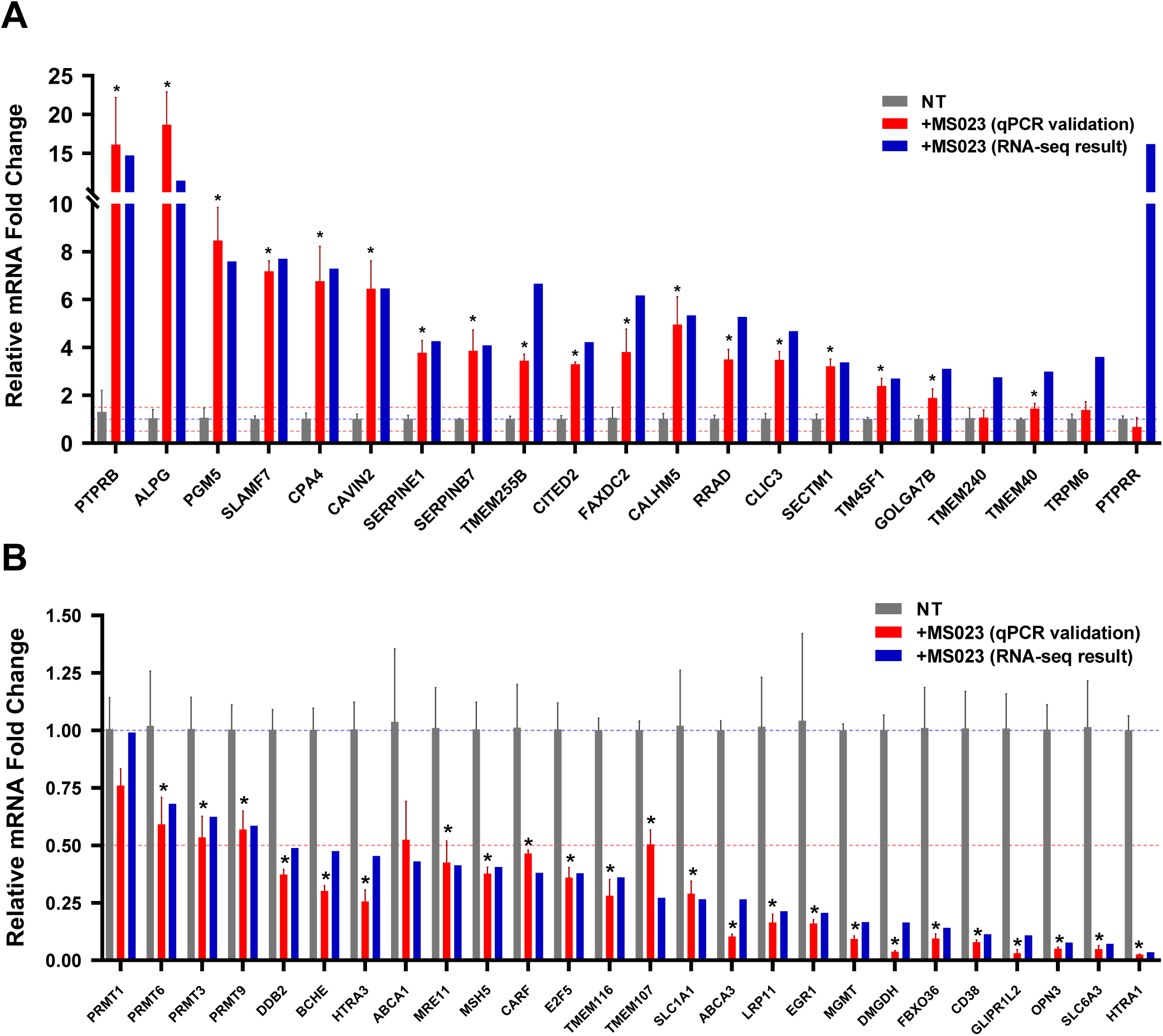
Validation of mRNA expression change of top selected up- and down-regulated genes. Top **(A)** up- and **(B)** down-regulated genes were picked, and expression changes were validated via RT-qPCR.

